# Neurocognitive mechanisms of social inferences in typical and autistic adolescents

**DOI:** 10.1101/850552

**Authors:** Gabriela Rosenblau, Christoph W. Korn, Abigail Dutton, Daeyeol Lee, Kevin A. Pelphrey

## Abstract

**Background:** Many of our efforts in social interactions are dedicated to learning about others. Adolescents with autism have core deficits in social learning, but a mechanistic understanding of these deficits and how they relate to neural development is lacking. The current study aimed to specify how adolescents with and with autism represent and acquire social knowledge and how these processes are implemented in neural activity.

**Methods:** Typically developing (TD) adolescents (N=26) and adolescents with autism (N=20) rated in the MR scanner how much three peers liked a variety of items and received trial-by-trial feedback about the peer’s actual preference ratings. In a separate study, we established the preferences of a new sample of adolescents (N=99), used to examine population preference structures. Using computational models, we tested whether participants in the MR study relied on preference structures during learning and how model predictions were implemented in brain activity.

**Results:** TD adolescents relied on average population preferences and prediction error (PE) updating. Importantly, PE updating was scaled by the similarity between items. In contrast, preferences of adolescents with autism were best described by a No-learning model that relied only on participants own preferences for each item. Model predictions were encoded in neural activity. TD adolescents encoded PEs in the putamen and adolescents with autism showed greater encoding of own preferences in the angular gyrus.

**Conclusions:** We specified how adolescents represent and update social knowledge during learning. Our findings indicate that adolescents with ASD rely only on their own preferences when making social inferences.

## Introduction

Social competences become increasingly important in adolescence – a transition period in which peers begin to outweigh family influence^1^. Despite the importance of peer socialization, adolescents make suboptimal social inferences, and in part this is likely due to ongoing development of brain regions^2–4^. Specifically, subcortical regions involved in emotion and reward processing, such as the ventral striatum, mature earlier than prefrontal regions, which support cognitive control and emotion regulation^5,6^. The medial prefrontal cortex (MPFC), a key region for learning from social feedback, is among the latest maturing brain regions^7,8^. In adolescents, the MPFC represents social predictions to a lesser degree than in adults, and that may account for a slower updating of social predictions through environmental feedback^9,10^.

Adolescents with Autism Spectrum Disorder (ASD) exhibit difficulties with social cognition^11,12^. Social attention and motivation accounts hold that reduced orientation to social stimuli early on in infants with ASD lead to cascading effects on social learning and social interaction later on^13–15^. With respect to adolescents with ASD, it is not clear whether social attention and motivation continue to change in this critical period for brain development. No study to date has investigated how social knowledge modulates updating in adolescents with ASD. As such, it remains unclear whether adolescents represent social knowledge when making social inferences and whether this knowledge guides learning. This is a particularly important question given that adolescents with ASD experience more negative social interactions than typical adolescents and an increased risk for comorbidities and emotional maladjustment^16,17^.

Computational modeling together with neuroimaging may provide a model for social inferences of adolescents with ASD and important insights into how such inferences are encoded in brain activity. The few studies that have described social decisions of adults with high autistic traits or ASD with computational models have found that these individuals represent others’ mental states less when making social inferences^18,19^. It remains unclear if this difference between typical and autistic individuals is accentuated in adolescence, a unique developmental period characterized by developmental asymmetries between social and general cognition ^20,21^.

In this study, we aimed to expand our previous approach of studying social learning in adolescence ^9^ by investigating how social knowledge structures shape learning. We investigated whether social inferences in typically developing (TD) adolescents and in adolescents with ASD rely on prior social knowledge of varying complexity. More specifically, we combined computational modeling with brain imaging to investigate how adolescents make inferences about what individuals from their peer group like and dislike. Participants received trial by trial feedback about the peer’s actual preference, which they could use to update inferences on similar subsequent items. Additionally, to assess social knowledge structures, we asked participants for their own preference after scanning. We also collected information about preference structures in a large, independent sample of adolescents. These metrics allowed us to compare how TD adolescents and adolescents with ASD use prior knowledge about themselves and others when making preference inferences.

In our previous report, we characterized social learning in typically developing adolescents and adults with computational models that assumed participants relied on prediction errors (PEs), the difference between initial ratings and subsequent feedback, and their own preferences to varying degrees. For this study, we constructed additional computational models that rely on peer preferences and scaling PEs based on similarities between preferences. We hypothesized that individuals with ASD would differ from typically developing (TD) adolescents in the extent to which they rely on prior knowledge about peers, i.e., represent peer preference structures, and in relying on social feedback to update predictions. We expected that differences in representing preference structures and prediction error encoding would scale with differences in neural encoding of social predictions in the MPFC.

## Materials and Methods

The study consisted of two assessments, an online preference survey in a large sample of adolescents to establish preferences of the adolescent population, and an fMRI experiment in a separate group of typically developing (TD) adolescents and adolescents with ASD. The experiments have been introduced in a previous report^9^ which also described part of the TD sample analyzed here. Briefly, the online survey contained pictures of activities, fashion and food items and a short demographic questionnaire. Survey participants rated how much they liked each item on a 10-point Likert scale ranging from 1 (not at all) to 10 (very much). The fMRI experiment was carried out using the same items. Participants were asked to infer the preferences of three people from their peer group and were presented with trial-by-trial feedback about the other’s actual preference rating. The rationale behind choosing multiple profiles was to increase the task’s ecological validity by increasing the generalizability of observed learning patterns to multiple distinguishable preference profiles. After the fMRI experiment, participants provided their own preferences for the items outside the scanner. The experimental procedures were conducted in compliance with the standards established by the universities’ Institutional Review Boards and the Declaration of Helsinki.

### Online survey participants

Adolescent participants (N=99; 55 females, mean age ± standard deviation = 15.7 ± 1.4 years; age range: 11-18 years) were recruited through flyers in the broader area of Washington DC and online advertisements. Participants and their parent/ guardians were invited to participate in the online preference study, which was available online through the Yale Qualtrics Survey Tool (yalesurvey.qualtrics.com). Participants were offered a $5 gift card upon completion (completion time was ~20 min).

### FMRI study participants

Twenty-four high-functioning adolescents with Autism Spectrum Disorder (ASD) were recruited from existing participant data bases at the Yale Child Study Center (12 female, mean age ± standard deviation = 14.8 ± 2.8 years; age range: 10-20 years). Prior to participation, diagnosis of all included individuals had been confirmed by expert clinicians using 1) the Autism Diagnostic Observation Schedule, Second Edition (ADOS-2; Lord et al., 2012) either Module 3 or Module 4 as deemed most appropriate by the assessing clinician, and 2) the Autism Diagnostic Interview-Revised (ADI-R, Lord et al., 1994). Four adolescents were excluded from the analysis due to excessive motion (absolute displacement mean relative displacement ≥ 1 mm, N=2) or because they interrupted the task / provided insufficient behavioral responses (we excluded participants with less than 50% of valid responses in any run, N=2) (see Table 1 for phenotypic characterization of the final sample).

**Table 1.**
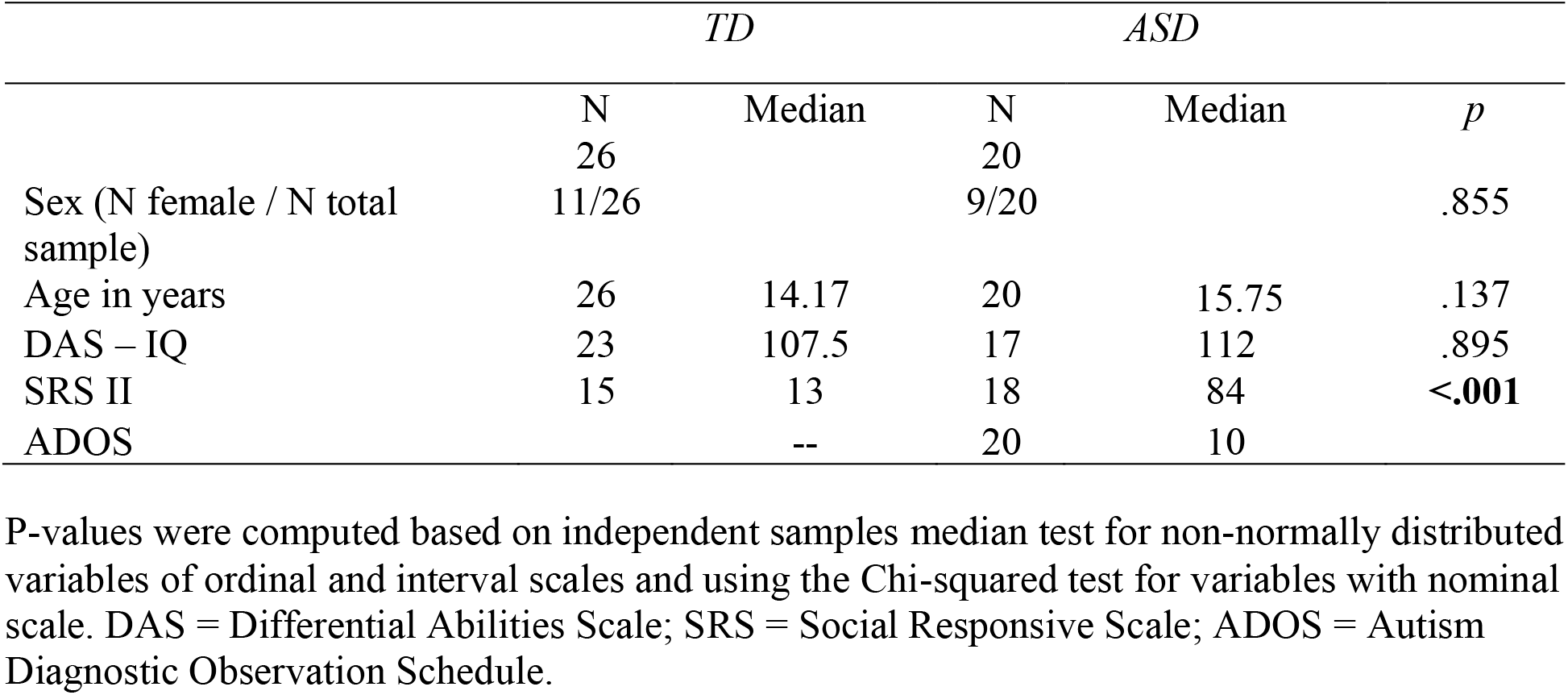
Demographics and symptom characteristics.

The ASD group was matched based on age, sex, and IQ with a sample of 26 typically developing (TD) adolescents (11 female, mean age ± standard deviation = 13.7 ± 2.5 years; age range: 9-18 years, Table 1). Note that data from most of this TD sample (N=24) has been included in a previous report^9^. Participants’ IQ was measured with the Differential Ability Scales (DAS, Elliott, 2012) by either clinically trained research staff (for TD participants) or licensed clinicians (for ASD participants).

Parents of participants in both groups provided an additional, independent assessment of social skills. They rated the social skills of their participating child on the Social Responsiveness Scale (SRS) first ^25^ or second ^26^ edition. For 4- to 18-year-olds, these two measures comprise the same items. The SRS measures the presence of impairments in reciprocal social behaviors, typically associated with Autism Spectrum Disorder (ASD). This measure has been validated as a measure of autistic traits in typical and ASD samples ^27^. TD adolescents in this sample all scored in the typical, non-autistic range (i.e., their total SRS T-score was below 59; mean ± standard deviation = 44.3 ± 4.5; range: 37-53). As expected, ASD participants scored significantly higher, mean ± standard deviation = 73.5 ± 13.9; range: 49-90).

### FMRI task design

Adolescent participants were asked to infer the preferences of three people from their peer group while performing three task runs in the scanner. Prior to each task run, participants received a brief introduction of the person whose preferences they had to infer on the respective run. Following their rating for an item, they received trial-by-trial feedback about the other’s actual preference rating (i.e., rating outcomes). Participants were instructed to get to know the persons during the task. Pictures of items were never repeated and participants were not given any strategies to make inferences or change them over the course of the task.

The task consisted of 120 items total (49 activity, 30 fashion, and 41 food items) including a relatively equal number of subcategory items (see supplemental Table S1 for a comprehensive list of stimuli and respective categories). These were divided into three equivalent item sets of 40 items each and assigned to one of three adolescent-preference profiles. Each profile was presented in a separate functional run (three runs in total). One run lasted for approx. 9 min 30 s (total task duration was ~30 min, total scan time was ~60 mina schematic depiction of it can be found in Fig.1).

**Fig. 1.**
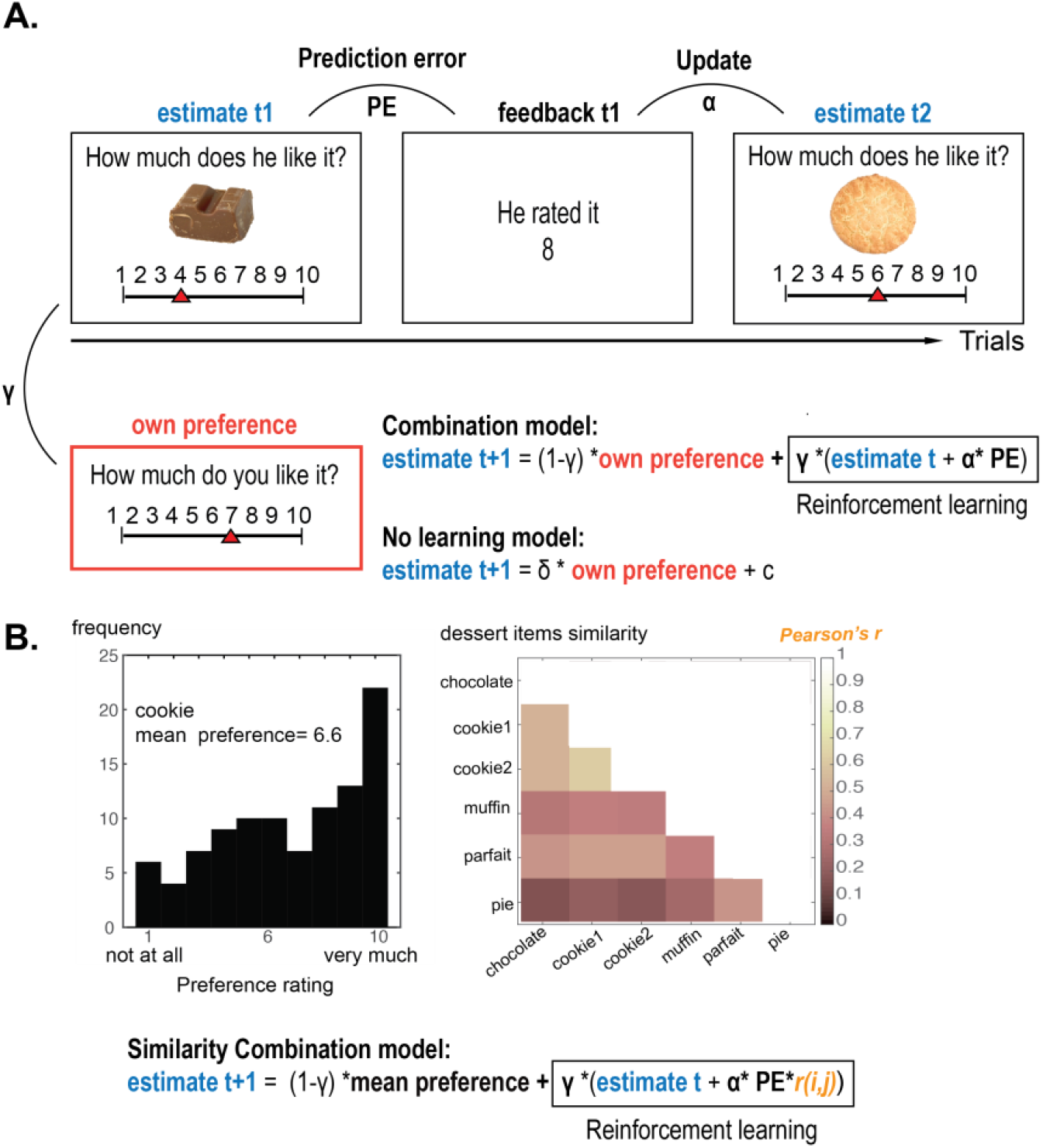
Experimental task and computational models. A. Example item for the preference task and computational models derived from task- and own preference information. B. Example distribution and mean rating for one item from the adolescent preference survey with N = 99 adolescents for the item cookies. Correlation matrices depict the relationship among ratings of participants’ self-preferences for different classes of items within an example sub-category (desserts). Mean preferences and preference similarity is used in the Similarity Combination model.

The overall frequency of missing responses did not differ between ASD and TD groups. (ASD: mean ± standard deviation = 6.5 ± 9.0; TD: mean ± standard deviation = 6.0 ± 8.0; *t* (44) = 1.38, *p* = .174).

After participants completed the fMRI task, they were asked to describe the individuals based on what they had learned from the task with open ended questions. We investigated whether ASD and TD participants formed similar social impressions of each person, by generalizing from the items presented to our pre-defined categories and even further to character traits. Two raters, unfamiliar with the objective of this study, coded the frequency of predefined classifications (i.e., categories and subcategories) and personality inferences in both ASD and TD groups (on average raters agreed on 79% of their classifications). We computed and compared between-group differences of predefined classifications and personality inferences, if both raters agreed.

Lastly, participants completed the same preference survey as the survey participants. They were asked to report on their own preferences for the fMRI task items (see Fig. 1 and the preference survey section for more details). Note that we assessed participants’ own preferences after the scanner task to avoid priming them towards their own preferences when judging those of their peers. Participants did not know beforehand that they would be asked for their own preferences in this study.

### Behavioral data analysis

Our main hypothesis is that learning strategies differ between TD and ASD groups. We hypothesized that TD adolescents would rely on sophisticated knowledge structures (i.e., preference similarities) and PEs, the difference between initial ratings and subsequent feedback, when making preference inferences about peers. We expected these differences in cognitive strategies to be reflected in the amount of PE reduction over the course of the task, whereby TD adolescents would reduce PEs over the course of the task more rapidly than adolescents with ASD. We also expected that learning strategies of TD adolescents would result in more holistic representations of preference profiles, leading to greater generalizations across items and item categories. These hypotheses were directly tested in model-free and model-based analyses, as detailed below.

To specify individual differences and the developmental trajectory of social learning in adolescents with ASD, we investigated linear and quadratic relationships between demographic variables (e.g., age as decimal values, autistic symptomatology) and PEs and model-based variables (e.g., learning rates from winning model). Given the growing literature on sex differences in ASD, sex was entered as a covariate in these analyses. We assessed whether variables were normally distributed using the Kolmogorov-Smirnov test. Non-parametric statistical tests were performed for non-normally distributed variables.

#### Model-free behavioral analysis

Participants could use the person’s feedback on previous items to inform preference inferences for upcoming items, thereby reducing PEs within item subcategories and beyond. For the model free analyses, we test participants’ overall accuracy in estimating the other persons by assessing PEs independent of the direction of deviation (positive or negative). We thus defined PEs as the absolute difference between participants’ ratings and the feedback they subsequently received. Note that for the model-based analysis described in the next section, PEs are used to adjust upcoming ratings up or down, and are therefore defined as the signed difference between rating and feedback.

To directly test the notion of generalized learning, we also asked participants to provide short descriptions of the persons after the scanner session. We expected TD adolescents to give more detailed descriptions of the persons, mentioning pre-defined categories and personality traits more frequently in their open answer descriptions of people after the task.

#### Computational modeling

To test whether learning strategies differ between ASD and TD participants, we devised and formally tested computational models that contain prior knowledge of varying complexity (i.e., own preferences, average peer preferences, preference similarities) and additional RL components (see the schematic depiction of main models in Fig. 1).

##### Model space

###### Main Models

Previously, our main model space consisted of three models, which assume participants adjust their estimated ratings of another person (ER)

1. by performing a simple linear transformation of their own preferences (OP) to predict the preferences of the other persons **(Model 1: No-learning)**;

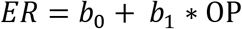
2. according to a variant of the Rescorla-Wagner reinforcement-learning (RL) rule (**Model 2: RL-ratings**). This model comprises the RL rule, which adjusts Expected Ratings of the other person’s preference *ER_t+1_* on the basis of the participant’s current estimates *ER_t_* and the Prediction Error *PE.* The PE is the difference between this current estimate and the current Feedback *F_t_*, which is the other person’s actual preference rating. The prediction error is weighted be the learning rate α, which is a free parameter.

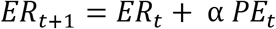

with

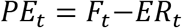
3. by using a weighted combination of RL and their own preferences to predict the others’ preferences (OP; **Model 3: Combination)**.

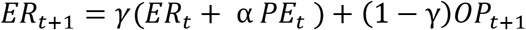

Here, we added four analogous models described below that additionally include simple and more sophisticated prior knowledge about preferences of peers. They scale prediction errors by preference similarities and / or substitute participants’ own preferences with average population preferences from the adolescent survey.

In **Model 4 (Simple knowledge)** participants are assumed to perform a linear transformation from population averages to participants’ ratings on the task. Participants base their estimated ratings (ER) of another person on the Mean Preference rating (MP) for the item in question.

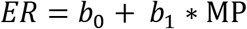

**Model 5 (Combination simple knowledge)** is analogous to the combination model (Model 3). Instead of own preferences this model contains a tradeoff between RL and the average population preferences MP.

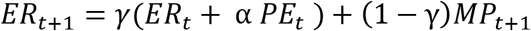

**Model 6 (Similarity-RL)** represents a more sophisticated version of the RL ratings (Model 2). An important difference between this and Model 2 is that for this model the integration of PEs is scaled by the preference similarity for the current item *i* and all subsequent items *i ∈ I* participants are going to see. The preference similarity is conceptualized as the correlation *r* between preferences of the independent population of 100 adolescents for the current item i and all subsequent items in the task set *I*.

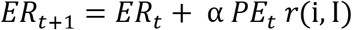

with

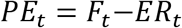

**Model 7 (Similarity-Combination)** represents a more sophisticated version of the Combination model (Model 3). Similar to the logic of the combination model, this model assumes participants’ ratings ER for the other person rely on a weighted combination of the RL rule with similarity weights (Model 5: RL-ratings) and average population preferences (*mean preferences MP*) to predict the others’ preferences.

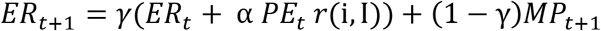

##### Testing initialization of estimated ratings

In accordance with our previous report, we initialized *ER_t_* of **Models 2, 3, 5-7** to the midpoint of the scale (5.5) for the first item of a new subcategory. We additionally tested whether estimating the initializing parameter resulted in better fit. This was not the case; models failed to converge for some participants and log-group Bayes factors (see explanation below) were higher than for the 5.5. initialization for the remaining subjects.

##### Model estimation and comparison

We used linear least squares estimation to determine best fitting model parameters. Optimization used a non-linear Nelder-Mead simplex search algorithm (implemented in the MATALB function *fminsearch*) to minimize the sum of squared errors of prediction (SSE) over all trials for each participant. We constrained parameter values to sensible *a priori* defined ranges (i.e., *α*, γ to a range of 0 to 1). For each model and each participant, we approximated model evidence by calculating the Bayesian Information Criterion (BIC), according to the following standard formula:

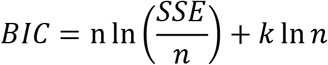

where *n* is the number of trials, and *k* the number of free parameters in the model. The latter is used to penalize model complexity. We report both fixed- and random-effects model comparison. For fixed-effects analyses, we computed log-group Bayes factors (BF) by summing BIC values for each tested model across participants and then subtracting the value of the reference model (**Model 2 RL-ratings**). According to the convention used here, smaller log-group BF indicate more evidence for the respective model versus the reference model. We furthermore investigated whether model frequencies for the winning models differ between ASD and TD groups. This random-effects family wise comparison was performed using the Bayesian Model Selection (BMS) procedure implemented in the MATLAB toolbox SPM12 (http://www.fil.ion.ucl.ac.uk/spm/; *spm_BMS,* Rigoux et al., 2014).

### fMRI data acquisition

Images were collected at the Yale University Magnetic Resonance Research Center on a Siemens 3T Tim Trio scanner equipped with a 12-channel head-coil. Whole-brain T1-weighted anatomical images were acquired using an MPRAGE sequence (repetition time [TR] = 2530 ms; echo time [TE] = 3.31 ms; flip angle = 7°; field of view [FOV] = 256 mm; image matrix 256 mm^2^; voxel size = 1 mm^3^; 176 slices). Field maps were acquired with a double echo gradient echo field map sequence, using 51 slices covering the whole head (TR = 731 ms; TE = 4.92 and 7.38 ms; flip angle = 50°; FOV = 210 mm; image matrix = 84 mm^2^; voxel size = 2.5 mm^3^). The experimental paradigm data were acquired in three runs of 285 volumes each (TR = 2000 ms; TE = 25 ms; flip angle = 60°; FOV = 220 mm; image matrix = 64 mm^2^; voxel size = 3.4 × 3.4 × 4 mm^3^; 34 slices).

### fMRI data analysis

#### Preprocessing and motion correction

FMRI data was processed using FEAT (FMRI Expert Analysis Tool) Version 6.00 of FSL. The first five volumes of each run were discarded to obtain a steady-state longitudinal magnetization. The *fsl_prepare_field_map* tool was used to correct for geometric distortions caused by susceptibility-induced field inhomogeneities. FSL’s *fsl_motion* outlier tool was used on non-motion-corrected functional data to detect timepoints corrupted by motion. A confound matrix was generated and used to regress out corrupt timepoints at the subsequent first-level analysis step. The inclusion requirement on this criterion was that no more than 20% of volumes could be identified as motion outliers; no data set exceeded this criterion. Further preprocessing steps included motion correction using MCFLIRT (Jenkinson et al. 2002), slice-timing correction, non-brain removal using BET (Smith 2002), spatial smoothing using a Gaussian kernel of FWHM 5 mm, and high-pass temporal filtering. Images were registered to the high resolution structural and to the Montreal Neurologic Institute (MNI) template using FLIRT (FMRIB’s Linear Registration Tool^29,30^). Furthermore, we tested whether groups differed with respect to the amount of motion during the experiment (i.e. mean relative displacement). Mean relative displacement across runs did not differ significantly between groups (Wilcoxon rank sum test: Z = 1.54, *p* = .124).

##### fMRI data analysis: Statistical model

Two general linear model (GLM) were set up for each participant and run to investigate how brain activity was modulated by model-free PEs (i.e., the differences between participants’ ratings and the received feedback) on a trial-by-trial basis (*model-free GLM*) and by variables derived from the winning behavioral model (*model-based GLM*). Both GLMs included two regressors for the distinct phases of the task: rating and feedback phases. The model-free GLM additionally included own preferences as parametric regressors in rating phases. Model free PEs were entered as parametric regressors in feedback phases. The parametric regressors of the model-based GLM were: model-predicted ratings, entered in rating phases, and model-predicted PEs, entered in feedback phases. In both GLMs, feedback numbers were entered as parametric regressors to account for variance explained by the simple presentation of numbers in the feedback phases. This additional regressor prevents erroneous assignment of variance associated with numbers only to the PE regressor. Finally, in both GLMs, nuisance regressors were included to account for subject motion: a confound matrix identifying outlier timepoints according to the DVARS metric ^31^ as well as six standard motion parameters. Correlations between parametric regressors for both groups were low (see Table 2).

**Table 2.**
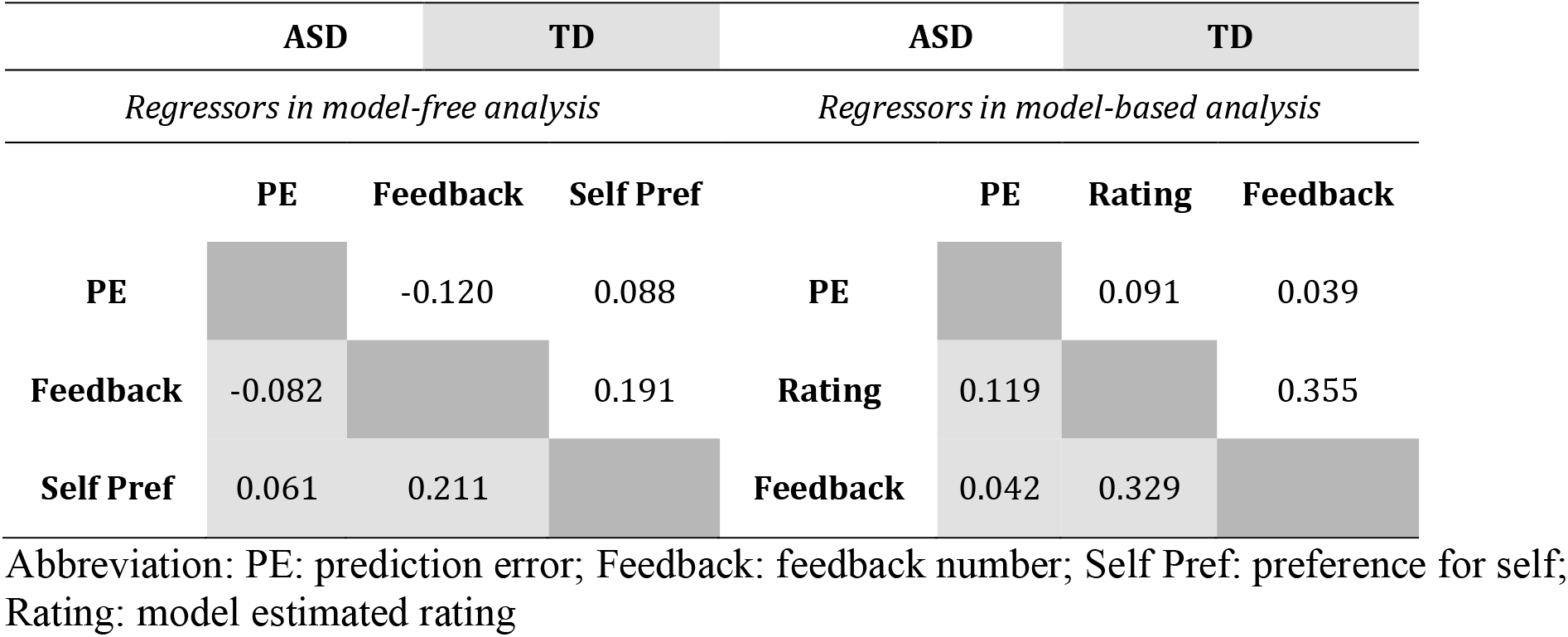
Correlations between parametric regressors in model-free and model-based analyses.

The runs were combined in a second-level within subject analysis and at the group level, we performed mixed-effects analyses on the contrast images using FLAME (FMRIB’s Local Analysis of Mixed Effects Stages 1 and 2 with automatic outlier detection and deweighting^32,33^). The model-free group analysis addressed the representation of model-free PEs in neural activity of TD and ASD groups as well as group differences (TD vs. ASD) in neural encoding of model-free PEs. The model-based group analysis tested the neural encoding of variables derived from the winning behavioral models (e.g., model derived ratings and PEs) in either group as well as differences between groups. All functional MRI analyses were family-wise cluster corrected at a z threshold of 2.3 and a p-threshold of p<0.001, in line with guidelines for adequate cluster correction with the FSL toolbox^34,35^.

## Results

### Differences in social inferences between typical and autistic adolescents

We tested group differences in task performance with two different metrics: unsigned model-free PEs (the numerical differences between estimates and feedback) and open-answer descriptions of preference profiles. Average accuracy that is average absolute model-free PEs did not differentiate between TD adolescents and adolescents with ASD. Median model-free PEs were on average equally high (Median for TD: 2.49; Median for ASD: 2.49; *H-test on median PEs*: *χ^2^*(45) = 0.07, *p*=0.790) and reduced over time to the same degree in both groups (t(44)=0.38, p=0.705). The relationship between average PEs and demographic variables, however, differed significantly between ASD and TD groups. In the ASD group, cognitive skills predicted performance on this social decision-making task. Adolescents with higher cognitive skills had lower average PEs (rho= −0.571, p=0.02). No significant relationship between IQ and PE magnitude was found in the TD group. Correlation coefficients differed significantly between groups (*Fisher’s r-to-z* = 2.82, *p* = 0.002).

Conversely, in the TD group, PEs were more related to age than in the ASD group (group difference in correlation coefficients: *Fisher’s r-to-z* = −2.97, *p* = 0.002), whereby a quadratic fit outperformed a linear fit (see supplemental Table S2). In line with reports of non-linear social development across adolescence^2^, younger and older TD adolescents had lower PEs than mid-adolescents.

Open-answer descriptions of peers in question revealed higher levels of generalization in the TD group. TD adolescents mentioned predefined categories (χ^2^(1) = 22.6, p < 0.001) and personality traits (χ^2^(1) = 3.52, p = 0.004) more often in their descriptions than adolescents with ASD. There were no group differences in the number of words participants wrote on average in each group (t(42) = 0.39, p = 0.696). Proportions of reported predefined categories and personality inferences for both groups are depicted in Fig. 2A.

**Fig. 2.**
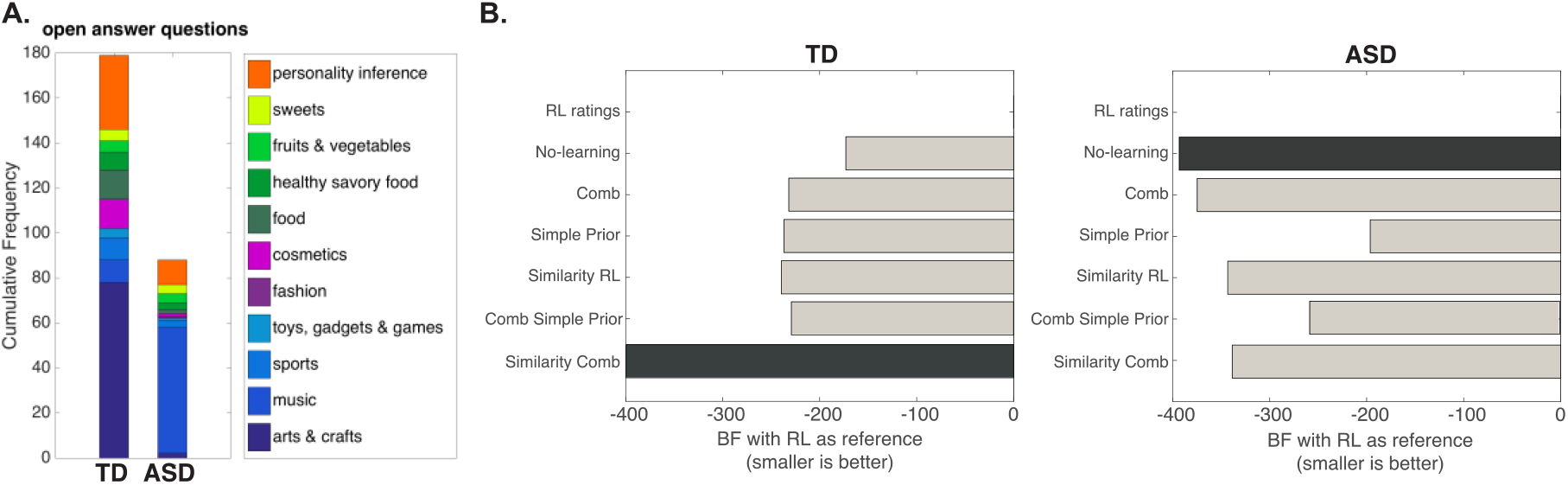
Group differences in social learning. A. Number of predefined categories and personality inferences mentioned in the open answer descriptions of TD and ASD adolescents. B. Bayes Factor (BF) representing model evidence for models with and without prior knowledge in TD and ASD groups. Preference ratings in the TD group rely on a combination of prediction error encoding scaled by similarity (the Similarity Combination model). Adolescents in the ASD group rely solely on their own preferences to rate those of others. The No-Learning model best explains ratings of adolescents with ASD. Abbreviations: ASD: Autism Spectrum Disorder, RL: Reinforcement learning, Comb: Combination, BF: Bayes Factor.

### Cognitive strategies underlying social inferences in typical and autistic adolescents

In this study, we extended the social inference model introduced previously^9^. In the previous study, we found that among the set of Models 1 to 3 the combination model including RL (Model 3) best described preference inferences of TD subjects. Here, we expanded our previous model space by models that additionally assume knowledge about peer preference structures. By comparing these models to previous models, we found that TD adolescents rely on knowledge about peer preferences. More specifically, preference predictions in the TD group were best captured by the Similarity-Combination model (Model 6). This model can be seen as a more fine-grained version of Model 3. Model 6 assumes that adolescents have initial knowledge about average peer preference and preference similarities for the item at hand and that they update this knowledge through RL. Feedback received about the peer in question generates a prediction error, which is scaled by item similarity; a large prediction error for an item is immediately applied to all items with a similar preference profile. In this way, the similarity matrix gradually changes to match the preference profile of the peer in question (see Fig. 3A for successive PE similarity updating).

**Fig. 3.**
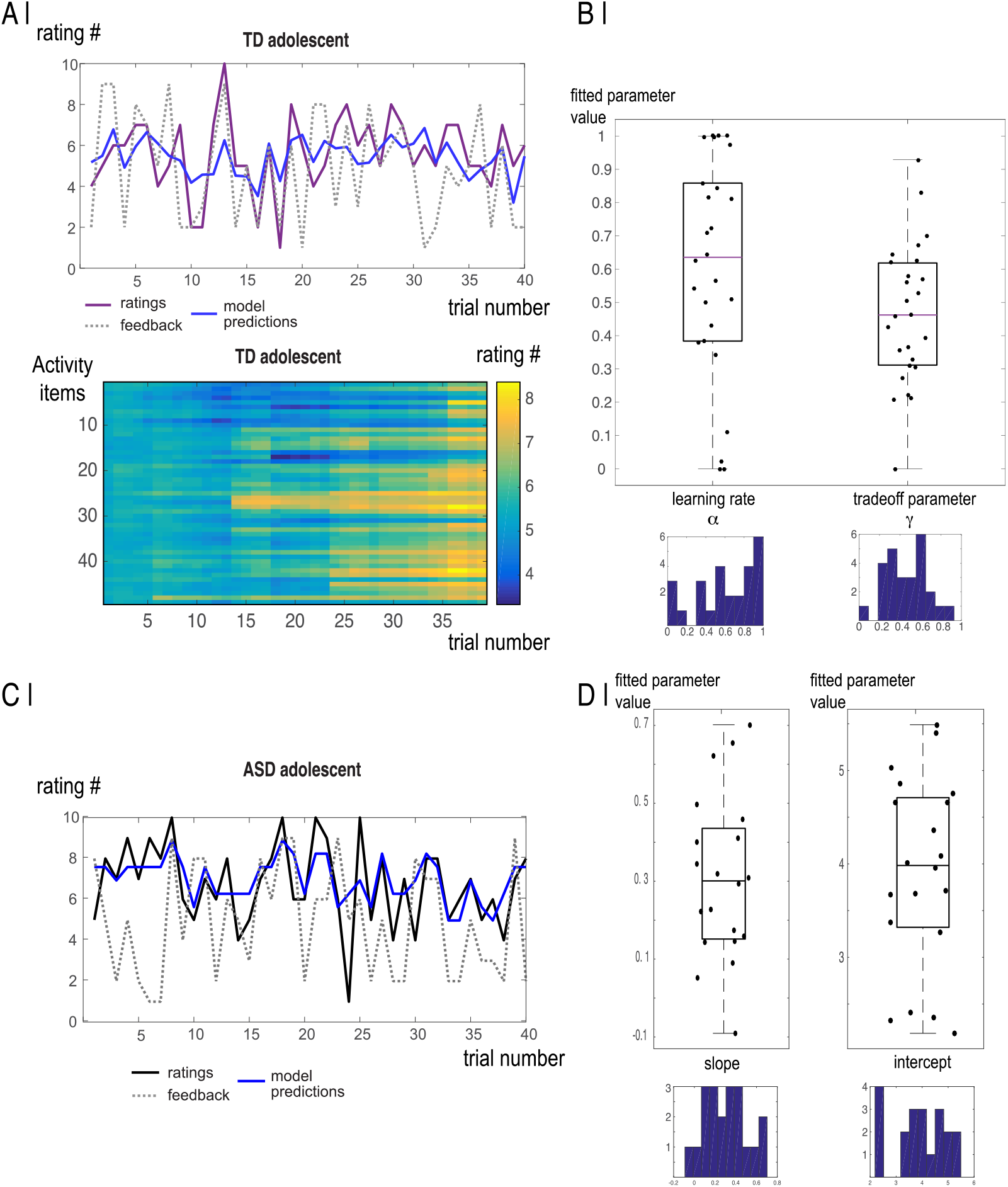
Estimates and fits of winning models in TD and ASD groups. A| Model fit and predictions in one TD adolescent during the task. Lower panel depicts trial by trial prediction error updating scaled by item similarity for all activity items simultaneously. At the beginning of the task estimates rely on item similarities between activity items only. Over the course of the task, participants use profile specific PEs to update estimates about activity items and scaled PEs by their similarity. B| Distributions of fitted parameter estimates from the Similarity-Combination model in TD adolescents. C| Model fit and predictions in one adolescent with ASD for the No-learning model that relies on own preferences only during the task. D| Distributions of parameter estimates from the No-learning model in adolescents with ASD. Abbreviations: TD: typically developing; ASD: Autism Spectrum Disorder

In general, in TD adolescents, models that incorporated prior knowledge about peer preferences outperformed those that relied on self-preferences as a reference. In contrast, the winning model in the ASD group was the No-Learning model (i.e. Model 2); the model solely relies on participants own preferences to predict those of others (Fig. 2B, Fig. 2C). Model comparison in each group suggested that the Similarity Combination was the best fitting model in the TD group and the No-learning model was the winning model in the ASD group (BF > 3). Parameters of the Similarity-Combination model were uncorrelated in the TD group, confirming the necessity of both parameters. The intercept and slope parameters for the No-learning model in the ASD group were, however, highly correlated (r=−0.911, p<0.001), the higher the intercept the lower the slope parameter (distributions of model parameters are depicted in Fig 3 B and D). We did not find conclusive evidence that groups use different models in a family wise random effects comparison (protected exceedance probability < .0.9). This is likely due to the great heterogeneity of ASD individuals.

#### Individual differences in model parameters

In line with previous reports of ongoing development in learning^9,36^, older TD adolescents with greater IQs had higher learning rates, meaning that they adjusted estimates more quickly based on task feedback (see supplemental Table S3). This relationship between model parameters and age, cognitive ability, and/or symptom severity was not significant in adolescents with ASD. Significant group differences were only found in the relationship between cognitive ability and parameter estimates (Fisher’s r-to-z=1.99, p=0.046).

### Neural activity scaled with task variables and model predictions

#### Model-free prediction errors

Trial-by-trial changes in model-free PEs scaled with activity in the MPFC in TD but not in ASD adolescents. The group difference, however, was not significant.

#### Model-based predictions

In line with our modeling results showing that adolescents with ASD rely on their own preferences when predicting preferences for others on the task. We find a stronger representation of own preferences in the angular gyrus extending into the precuneus cortex of ASD compared to TD adolescents during rating phases. Similarly, model-predictions derived from the winning computational model of TD adolescents, were encoded in brain activity during the task. Specifically, model-derived PEs from the winning Similarity Combination model were represented in the right caudate and putamen of TD adolescents during feedback phases (see Fig. 4, supplemental Tables S4 and S5).

**Fig. 4.**
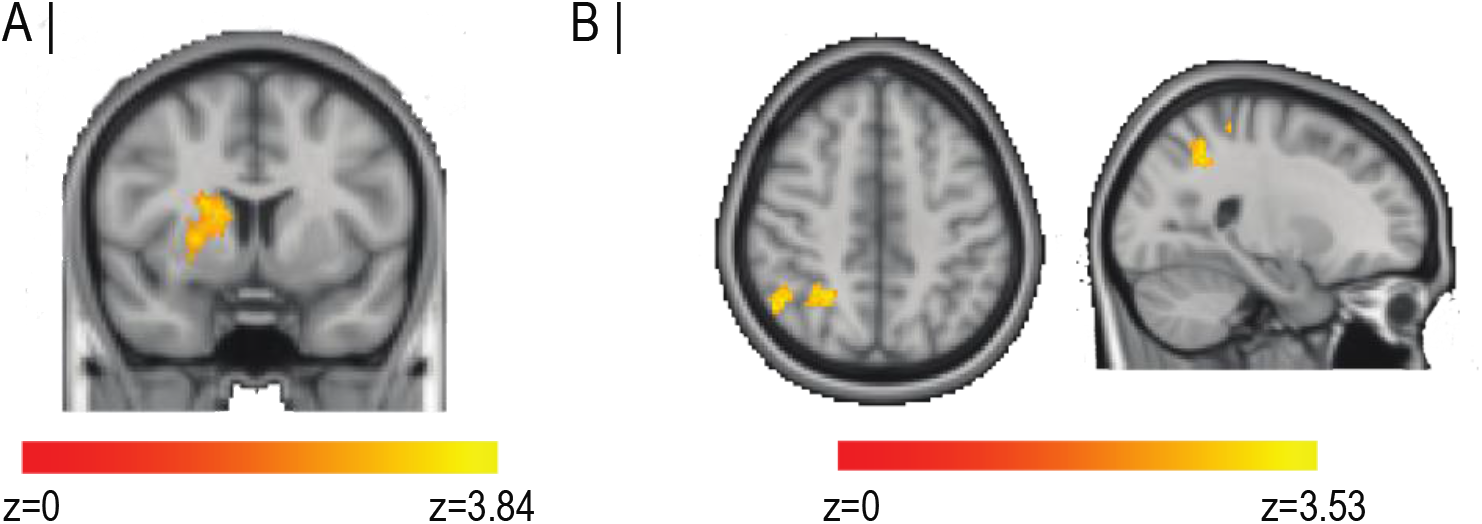
Parametric modulation of brain activity during the social inference task. A| Brain activity scales with PEs estimated by the Similarity-Combination model in typically developing adolescents. B| Brain activity that correlates more strongly with own preferences in the ASD vs. TD group.

## Discussion

This study investigated how adolescents with and without autism build knowledge representations of their peers; we tested whether adolescents rely on preexisting social knowledge to make preference inferences and how their knowledge is updated during learning about a specific peer. Based on preference profiles from an independent sample of adolescents, we devised computational models that combine knowledge about peers at varying levels of complexity with PE updating. TD adolescents relied on a combination of average population preferences and PE updating scaled by the similarity between current and upcoming items to optimize preferences of peers. In the ASD group, a simple No-learning model that only relies on participants’ own preferences provided the best fit to the data. Predictions of the winning computational models were corroborated by our neuroimaging results. In the TD group, model-derived PEs scaled with brain activity in the putamen. There was also a group difference in the extent to which own preferences were encoded in brain activity. In ASD adolescents, own preference ratings were more closely related to activity in the angular gyrus extending into the precuneus than in TD adolescents.

### Social learning through prediction errors

TD and ASD adolescents were asked to infer preferences of peers and could update these predictions based on trial-by-trial feedback. TD adolescents were able to build more generalized representations of the peers in question compared to ASD adolescents. In open ended questions after the experiment, participants generalized from information about specific items to broader item categories and personality inferences. Our results are in line with a vast body of literature showing that individuals with ASD have core deficits extracting general mental state inferences from social observations^37–39^. TD and ASD participants did not differ in the magnitude of PEs or in PE changes over time. The relationship between PEs and demographic variables, however, differed between groups. PEs were significantly related to age in the TD group – with mid adolescents having larger PEs than young and late adolescents. This non-linear trajectory of social development is a well replicated finding. Adolescence is a unique period for social development; mid-adolescents have higher prediction errors^40^ and lower learning rates than younger and older peers, which means that they learn more slowly from social feedback^9,41^. We did not find such ongoing social development in adolescents with ASD. Instead, ASD adolescents with higher IQs had overall lower PEs on the social inference task. Previous studies also found that language and academic skills improve social adjustment of adolescents with autism^42,43^, while for TD adolescents, social and cognitive skills are relatively independent from each^44,45^.

### Social knowledge representation and updating

The main aim of this study was to expand our previous computational modeling approach by investigating how social knowledge structures shape learning. Theories about knowledge acquisition postulate that humans acquire and represent abstract knowledge unconsciously and automatically by extracting statistical regularities from situations based on a set of rules^46–48^. Previous studies have shown that such knowledge representations are activated during learning^49,50^. Our study makes an important contribution to this literature by specifying how social knowledge structures are represented and updated during learning through task-based feedback. The computational modeling approach, revealed that choices of TD adolescents rely on average peer preferences and PEs scaled by preference similarities. If, for instance, a participant previously learned about their peer’s preference for bananas and now has to judge how much the peer likes apples, the participant would rely on the average population preference for apples and on how much the peer liked bananas before, given the degree to which preferences for bananas and apples are related.

In contrast, the model that best described inferences of adolescents with ASD was a simple No-learning model that just relies on participants’ preferences for the item at hand (an inference about how much the peer likes apples, relies on how much the participant likes apples). This model differs from the model of TD adolescents in two crucial ways: first, it does not rely on preexisting knowledge about population preferences and second, it does not rely on PE updating from feedback about the person’s actual preferences. A failure to encode social PEs and adjust ratings respectively may account for the observed social rigidity of individuals with ASD^51^ and explain their lack of preexisting social knowledge structures^52^. Due to high variability of model fits in the ASD group, we could not firmly establish that the winning computational models differed between groups. The fact that a No-learning strategy provided the best fit for the ASD group is in line with two recent accounts of ASD, which posit that individuals with ASD experience deficits in calculating decision variables and/or representing the statistical structure of the environment^19,51–54^. When making social inferences, adolescents with ASD used less informative knowledge (i.e., themselves versus population preferences) and did not include an information updating mechanism based on environmental feedback.

### Neural mechanisms underlying social inferences

The computational models that best described participants’ behavior were validated by our neuroimaging results. In typically developing adolescents, PEs estimated by the winning model were encoded in the putamen and caudate, parts of the striatum that are critically involved in decision making and learning from PEs^55–58^. Adolescents show increased activity of these regions when computing decision values and this has been typically linked to heightened dopaminergic PE responsivity and to increased reward-seeking tendencies of adolescents^21,56^. Model free PEs scaled with activity of the MPFC, a region that has been extensively implicated in encoding and updating others’ mental states^59–61^. The discrepancy between regions encoding model-based and model-free PEs highlights the difference between these decision variables: Model-free errors are conceptualized as the discrepancy or absolute differences between participants’ ratings and the subsequent feedback. Model-based PEs are signed error values that adjust model estimates up and down. Model estimates are not the actual rating of the participant but reflect population means and the similarity structure used by the model. Dopaminergic neurons in the ventral striatum may be sensitive to this signed error term, which contains the valence of the preference information (peer likes something more or less than predicted).

Unlike TD adolescents that encoded both model-based and model-free PEs in neural activity, we did not find neural encoding of PEs in adolescents with ASD. ASD participants encoded own preferences more strongly in the angular gyrus (AG) extending into the precuneus cortex compared to TD adolescents, corroborating their reliance on own preferences when making inferences about peers during the task. The AG and precuneus have been identified as integrative regions that store and represents conceptual knowledge and support attention to relevant information^62,63^. The precuneus, in particular, has been repeatedly linked to social deficits of individuals with ASD^64,65^ and changes in precuneus activity scale with improvements in social cognition^66^. To firmly establish the roles of these regions in encoding decision variables during social learning of individuals with ASD, this finding has to be replicated in future studies with larger sample sizes. In a similar vein, the current study lacks a representative sample of ASD preference profiles to rule out that ASD adolescents rely on representations about peers with ASD. However, we did not actually tell any of the participants whether the preferences came from adolescents with or without autism. Further, it is questionable that adolescents with ASD have the opportunity to build elaborate social knowledge structures about peers with ASD, and, if different, these knowledge structures would likely be suboptimal for predicting preferences of peers without ASD.

In summary, our study sheds light on adolescent social development in several important ways: first, it extends our previous account of social learning in adolescence by showing that typical adolescents represent sophisticated social knowledge structures during learning and that these structures are continuously updated through input from the environment. Second, to our knowledge, this is the first study that specifies the cognitive strategies underlying social learning about others’ preferences in autism. Finally, our results provide evidence for the usefulness of a neuro-computational approach in describing ongoing social development and social deficits of individuals with neuropsychiatric disorders

