## supplemental tables for "Neurocognitive mechanisms of social inferences in typical and autistic adolescents"

**Table S1 Items depicted in the preference task**

| 1 | accordion 01 | 46 | chocolate |
| --- | --- | --- | --- |
| 2 | accordion 02 | 47 | cookie 01 |
| 3 | apple 01 | 48 | cookie 02 |
| 4 | apple 02 | 49 | cube |
| 5 | asian food 01 | 50 | cucumber |
| 6 | asian food 02 | 51 | dice |
| 7 | asian food 03 | 52 | dominos |
| 8 | asian food 04 | 53 | drums |
| 9 | bacon on toast | 54 | earrings |
| 10 | college bag 01 | 55 | fitness 01 |
| 11 | college bag 02 | 56 | fitness 02 |
| 12 | girly bag | 57 | fitness 03 |
| 13 | shopping bag | 58 | frisbee 01 |
| 14 | women's bag | 59 | frisbee 02 |
| 15 | bagel | 60 | gratin |
| 16 | ball | 61 | greens 01 |
| 17 | ballet shoes | 62 | greens 02 |
| 18 | beads 01 | 63 | guitar 01 |
| 19 | beads 02 | 64 | guitar 02 |
| 20 | berries | 65 | head band |
| 21 | bike 01 | 66 | headphones 01 |
| 22 | bike 02 | 67 | headphones 02 |
| 23 | bike 03 | 68 | hummus |
| 24 | binocular | 69 | ice skates 01 |
| 25 | biscuits | 70 | ice skates 02 |
| 26 | boomerang 01 | 71 | kiwi |
| 27 | boomerang 02 | 72 | lip stick |
| 28 | bowling | 73 | makeup |
| 29 | bracelet | 74 | makeup |
| 30 | bun | 75 | microscope |
| 31 | burger 01 | 76 | mikado |
| 32 | burger 02 | 77 | mixed fruit |
| 33 | cake 01 | 78 | muffin |
| 34 | cake 02 | 79 | nail polish 01 |
| 35 | camcorder | 80 | nail polish 02 |
| 36 | camera 01 | 81 | nail polish 03 |
| 37 | camera 02 | 82 | necklace 01 |
| 38 | candy 01 | 83 | necklace 02 |
| 39 | candy 02 | 84 | notebook 01 |
| 40 | cards | 85 | notebook 02 |
| 41 | cherries | 86 | oil colors 01 |
| 42 | chess | 87 | oil colors 02 |
| 43 | chicken | 88 | orange |
| 44 | chips 01 | 89 | paper |
| 45 | chips 02 | 90 | parfait |
| 91 | pencil | 103 | shaving brush |
| 92 | piano | 104 | shaving brush |
| 93 | pie | 105 | skateboard |
| 94 | pizza | 106 | sneaker |
| 95 | radio 01 | 107 | sneaker |
| 96 | radio 02 | 114 | snorkeling gear |
| 97 | ribbon | 115 | soup |
| 98 | ring | 116 | sports bag 01 |
| 99 | roller skates | 117 | sports bag 02 |
| 100 | rucksack 01 | 118 | sunglasses 01 |
| 101 | rucksack 02 | 119 | sunglasses 02 |
| 102 | sandwich | 120 | taco |

**Table S2 Hierarchical regression testing the linear and non-linear relationships of PE a in the TD adolescent group**

|  | Unstandardized Coefficients | | |  | | | |
| --- | --- | --- | --- | --- | --- | --- | --- |
| Significant model^1^ | *B* | *SE* | *t* | | *F_1,21_* | *p* | *r^2^* |
| age squared | -.003 | 8.4*10^-4^ | -3.22 | | 10.37 | .004 | .257 |

^1^Regression tested the effect of IQ, linear and nonlinear age effects (age squared). TD = typically developing

**Table S3 Hierarchical regression testing the linear and non-linear relationships of learning rates estimated with the Similarity Combination model in the TD adolescent group**

|  | Unstandardized Coefficients | | |  | | | |
| --- | --- | --- | --- | --- | --- | --- | --- |
| Significant model^1^ | *B* | *SE* | *t* | | *F_1,19_* | *p* | *r^2^* |
| Intercept | -1.815 | .747 | -2.43 | |  | 0.03 |  |
| IQ | .013 | .004 | 2.79 | | 4.911 | .011 |  |
| age | .066 | .03 | 2.48 | | 4.911 | .022 | .2939 |

^1^Regression tested the effect of IQ, linear and nonlinear age effects (age squared). TD = typically developing

**Table S4. Regions showing parametric modulation by model-free prediction errors. Clusters are whole-brain FWE corrected for multiple comparisons at p <0.001 with a cluster-defining threshold of z=2.3.**

|  | Side | Peak voxel MNI coordinates (mm) | | | Cluster size (Voxel) | Peak z score |
| --- | --- | --- | --- | --- | --- | --- |
|  |  | x | y | z |  |  |
| Regions showing positive trial-by-trial correlation with PE during feedback phase | | | | | | |
| *TD Adolescents* |  |  |  |  |  |  |
| *Cluster 1* |  |  |  |  | *812* |  |
| Frontal Pole |  | 22 | 48 | 30 |  | 3.5 |
|  |  | 10 | 48 | 42 |  | 3.39 |
|  |  | 26 | 34 | 24 |  | 3.39 |
| Medial Prefrontal Cortex |  | 18 | 34 | 23 |  | 3.35 |
|  |  | 16 | 38 | 23 |  | 3.32 |
|  |  | 10 | 54 | 14 |  | 3.14 |
| *ASD Adolescents* |  |  |  |  |  |  |
| No Significant Clusters |  |  |  |  |  |  |
| *ASD>TD* |  |  |  |  |  |  |
| No Significant Clusters |  |  |  |  |  |  |
| *TD>ASD* |  |  |  |  |  |  |
| No Significant Clusters |  |  |  |  |  |  |

**Table S5. Regions showing parametric modulation by variables derived from winning computational models. Clusters are whole-brain FWE corrected for multiple comparisons at p <0.001 with a cluster-defining threshold of z=2.3.**

|  | Side | Peak voxel MNI coordinates (mm) | Cluster size (Voxel) | Peak z score |
| --- | --- | --- | --- | --- |

| Regions showing positive trial-by-trial correlation own preferences during rating phase | | | | | | |
| --- | --- | --- | --- | --- | --- | --- |
| *TD Adolescents* |  |  |  |  |  |  |
| No Significant Clusters |  |  |  |  |  |  |
| *ASD Adolescents* |  |  |  |  |  |  |
| No Significant Clusters |  |  |  |  |  |  |
| *ASD>TD* |  |  |  |  |  |  |
| Cluster 1 |  |  |  |  | 942 |  |
| Angular gyrus | R | 24 | -54 | 42 |  | 3.53 |
|  |  | 42 | -46 | 56 |  | 3.49 |
|  |  | 32 | -52 | 50 |  | 3.46 |
|  |  | 28 | -56 | 50 |  | 3.45 |
|  |  | 46 | -46 | 56 |  | 3.48 |
|  |  | 46 | -54 | 44 |  | 3.45 |
| *TD>ASD* |  |  |  |  |  |  |
| No Significant Clusters |  |  |  |  |  |  |
| Regions showing positive trial-by-trial correlation with PEs derived from Model 6 (feedback phase) | | | | | | |
| *TD Adolescents* |  |  |  |  |  |  |
| Cluster 1 |  |  |  |  | 742 |  |
| Caudate | R | 22 | 10 | 14 |  | 3.84 |
|  |  | 18 | 12 | 10 |  | 3.42 |
| Putamen | R | 24 | 6 | 2 |  | 3.31 |
|  |  | 22 | 0 | 10 |  | 3.26 |
| *ASD Adolescents* |  |  |  |  |  |  |
| No Significant Clusters |  |  |  |  |  |  |
| *TD >ASD* |  |  |  |  |  |  |
| No Significant Clusters |  |  |  |  |  |  |
| *ASD > TD* |  |  |  |  |  |  |
| No Significant Clusters |  |  |  |  |  |  |
